# TRONCO: an R package for the inference of cancer progression models from heterogeneous genomic data

**DOI:** 10.1101/027474

**Authors:** Luca De Sano, Giulio Caravagna, Daniele Ramazzotti, Alex Graudenzi, Giancarlo Mauri, Bud Mishra, Marco Antoniotti

**Affiliations:** Department of Informatics, Systems and Communication, University of Milano-Bicocca, Milan, Italy; School of Informatics, University of Edinburgh, Edinburgh, UK; Institute of Molecular Bioimaging and Physiology of the Italian National Research Council (IBFM-CNR), Milan, Italy; Courant Institute of Mathematical Sciences, New York University, New York, USA; Milan Center for Neuroscience, University of Milan-Bicocca, Italy

## Abstract

**Motivation:** We introduce `TRONCO` (TRanslational ONCOlogy), an open-source R package that implements the state-of-the-art algorithms for the inference of cancer progression models from (epi)genomic mutational profiles. `TRONCO` can be used to extract population-level models describing the trends of accumulation of alterations in a cohort of cross-sectional samples, e.g., retrieved from publicly available databases, and individual-level models that reveal the clonal evolutionary history in single cancer patients, when multiple samples, e.g., multiple biopsies or single-cell sequencing data, are available. The resulting models can provide key hints in uncovering the evolutionary trajectories of cancer, especially for precision medicine or personalized therapy.

**Availability:** `TRONCO` is released under the GPL license, it is hosted in the Software section at http://bimib.disco.unimib.it/ and archived also at bioconductor.org.

## 1 Introduction

Cancer develops through the successive expansions of clones, in which certain (epi)genomic alterations, called drivers, confer a fitness advantage and progressively accumulate, in a context of overall scarcity of resources [6]. Specifically, in Nowell’s seminal work, tumor evolution is described in terms of stepwise genetic variation such that growth advantage is the key for the survival and prorogation of the clones. Therefore, one can define *cancer progression models*, in terms of probabilistic *causal graphical models*, where the conditional dependencies and the temporal ordering among these alterations are described, revealing the evolutionary trajectories of cancer at the (epi)genome level.

We further distinguish [3]. (*i*) *ensemble-level* progression models, describing the statistical trends of accumulation of genomic alterations in a cohort of distinct cancer patients. Such models describe the temporal partial orders of fixation and accumulation of such alterations and represent population-level trends; and (*ii*) *individual-level* models, thus accounting for the specific evolutionary history of cancer clones in individual tumors. Such models thus impute the ancestry relations of the observed clones.

Even if the inference of such models is further complicated by a series of theoretical and technical hurdles, such as, e.g., *intra*- and *inter-tumor* heterogeneity and the effective detection of drivers, it can benefit from the increasing amount of *next-generation sequencing* (NGS) data, currently available through public projects such as The Cancer Genome Atlas (TCGA, https://tcga-data.nci.nih.gov). Usually, such databases provide *cross-sectional* (epi)genomic profiles retrieved from single biopsies of cancer patients, which can be used to extract ensemble-level models; but higher resolution data such as *multiple-biopsies*, or even *single-cell* sequencing data are becoming more accessible and reliable, which can be used to infer individual-level models.

Here we introduce TRONCO (TRanslational ONCOlogy), an R package built to infer cancer progression models from heterogeneous genomic data (in the form of alterations persistently present along tumor evolution.) Currently, TRONCO provides the implementation of two algorithms: (*i*) CAPRESE (Cancer PRogression Extraction with Single Edges [7]), and (*ii*) CAPRI (CAncer PRogression Inference [8]), both based on Suppes’ theory of *probabilistic causation* [9], but with distinct goals and properties (see Software Implementation).

TRONCO, in its current form and perspective, should be thought of as a tool that provides the implementation of up-to-date solutions to the progression inference problem. At the time of the writing it can be effectively used as the final stage of a modular pipeline for the extraction of ensemble-level cancer progression models from cross-sectional data [3]. In such a pipeline input data are pre-processed to (*i*) stratify samples in tumor subtypes, (*ii*) select driver alterations and (*iii*) identify groups of fitness-equivalent (i.e., mutually exclusive) alterations, prior to the application of the CAPRI algorithm. The resulting ensemble-level progression models depict the evolutionary dynamics of cancer, with translational impacts on diagnostic and therapeutic processes, especially in regard to precision medicine and personalized drug development.

From the complementary perspective, TRONCO can also exploit the CAPRESE algorithm to infer the clonal evolutionary history in single patients when multiple samples are available, as in the case of multiple biopsies and/or single-cell sequencing data, as long as the set of driver events is selected; see [3].

## 2 Software Implementation

TRONCO implements a set of R functions to aid the user to extract a cancer progression model from genomic data. At a high-level, these function shall help to import, visualize and manipulate genomic profiles – regardless of their source – eventually allowing the implemented algorithms to run and assess the confidence in a model.

The basics steps of TRONCO’s usage are shown in Figure 1. In panel (a) we show multiple input alterations (e.g., somatic mutations or copy number alterations) either from a cohort of patients, or a unique patient (e.g., multi-region or single-cell sequencing); in panel (b) we show an oncoprint visualization from the tool, i.e., a matrix whose columns represent samples and rows the alterations and their presence per sample; panel (c) shows an inferred graphical Bayesian progression model obtained with one of the available algorithms; finally, in panel (d) we show data supported for processing in the tool. For a more detailed explanation of the implementation of the package see [1].

**Figure 1.**
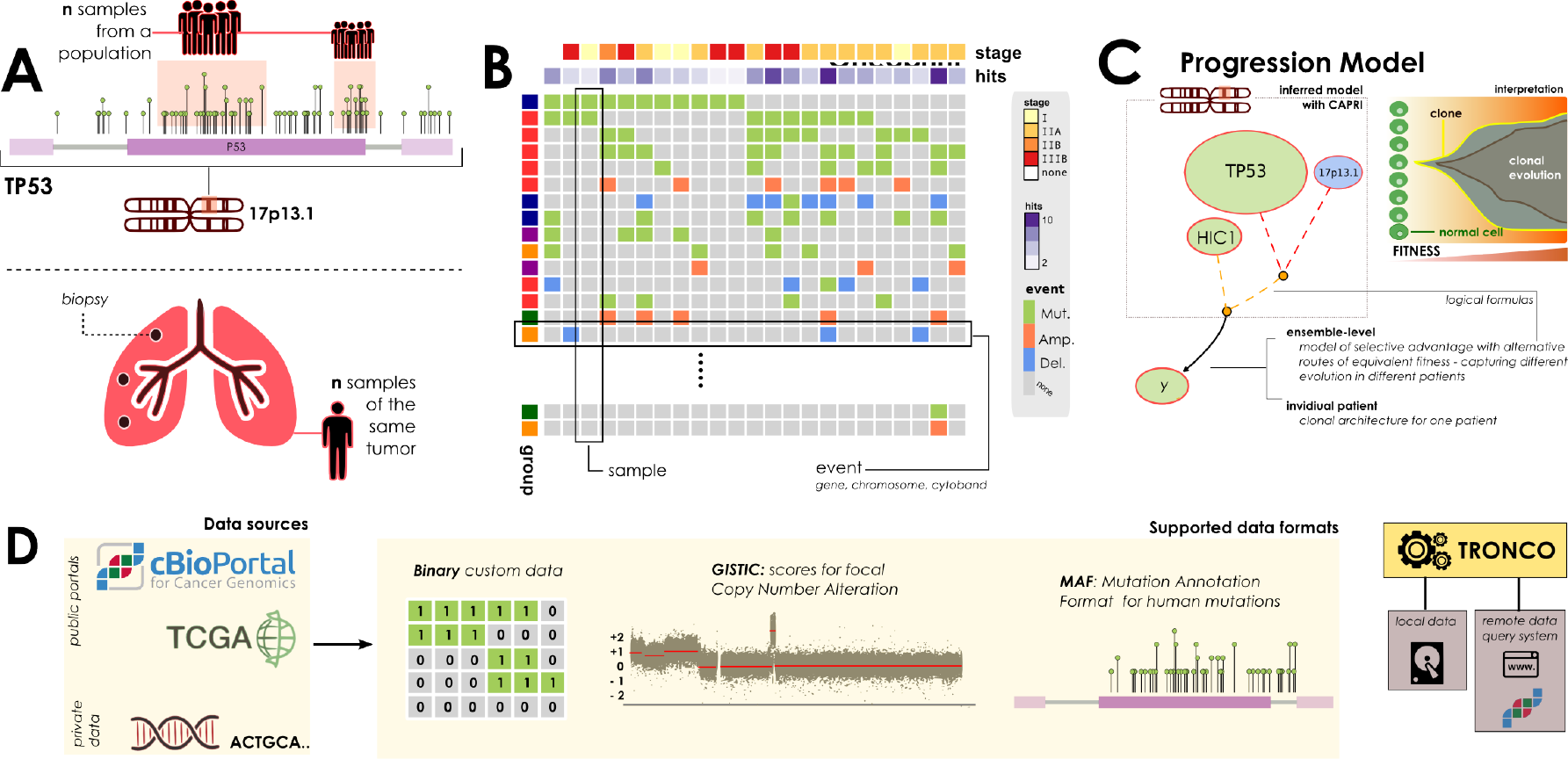
(A) TRONCO can process either alterations (e.g., somatic mutations or wider chromosomal lesions) in a cohort of independent samples (top lolliplot diagram), or a set of multiple snapshots from a unique patient (e.g., multi-region or single-cell, bottom panel). (B) Oncoprints allow the user to visualize the data that the tool is processing. Regardless of the source, each row represents a certain alteration - at a custom resolution depending on the cancer under study - and each column a sample. (C) A model inferred with the tool might outline cancer evolution occurring in a population ensemble or in an individual patient. Graphically, alterations are represented as nodes with different colors (e.g., green mutations and blue homozygous deletions). Algorithms such as CAPRI allow describing alterations with logical formulas, in an attempt to find their role as a “group” (see [8] for details); we picture such groups with dashed lines. In the panel, we show a hypothetical ensemble-level model predicting a selection pressure on two genes mapped to 17p13, tp53 and hic1, as it may be inferred by analyzing samples harbouring either tp53/hic1 mutations or homozygous deletions in the cytoband where any of these two genes map, i.e., here for purely explanatory cases we suppose just tp53, which maps to 17p13.1. The model suggests a trend of selection towards mutations in gene y, which shall be interpreted as a set of preferential clonal expansions characteristic of the population of analyzed samples, involving alterations of the functions mapped to 17p13 and y. (D) TRONCO supports three data types. Custom data, which is supposed to be provided as a *binary input matrix* storing the presence (1) or absence (0) of a certain alteration in a sample. Or, standard data formats such as the *Mutation Annotation Format* (MAF) for somatic mutations, as well as the *Genomic Identification of Significant Targets in Cancer* (GISTIC) format for focal Copy Number Variations. Data can be generated by custom experiments, or collected - along with other “omics” - from public databases such as TCGA and cBio portal. For the latter, cBio portal, TRONCO implements a query system to fetch data with minimal effort. The tool engine can then be used to manipulate genomic profiles – regardless of their source – and run progression inference algorithms.

**Data loading and manipulation**. Common formats to store data used to extract progression models can be natively imported. These include, for instance, the *Mutation Annotation Format* (MAF) for somatic mutations, as well as the *Genomic Identification of Significant Targets in Cancer* (GISTIC) format to store focal Copy Number Variations. The tool can exploit the cBio portal for Cancer Genomics, which collects among others TCGA projects, to access freely available instances of such data [4].

TRONCO provides functions for data preprocessing to, e.g., select a certain subset of alterations, or samples or any abstraction which might be appropriate according to the cancer being studied.

**Visaualization and interaction with other tools**. TRONCO implements an *oncoprint* system to visualize the processed data. Datasets can be exported for processing by other tools used to, e.g., stratify input samples and detect groups of mutually exclusive alterations, which include the *Network Based Stratification* [5] and *MUTEX* [2] tools. TRONCO allows the visualization of the inferred models.

**Model inference and confidence**. TRONCO provides two algorithms: (*i*) CAPRESE, which uses a *shrinkage*-like estimator to infer *tree*-models of progression, and (*ii*) CAPRI, which extracts more general *direct acyclic graphs* (DAG) - thus allowing for confluent evolution and complex hypothesis testing – by combining *bootstrap* and *maximum likelihood estimation*. CAPRESE and CAPRI both rely on the same theory of *probabilistic causation*, but with distinct goals and properties. The former reconstructs tree models of progressions, while the latter general directed acyclic graphs. Both methods are agnostic to the type of input data (i.e., whether its an ensemble or an individual tumor), but shall be used in different contexts as they produce different types of models. Indeed, CAPRESE is better at extracting cancer evolution in a single individual as in that case trees capture branched evolution and trunk events, which shall suffice to describe clonal evolution. Instead, when heterogeneity might result in multiple evolutionary routes with common downstream alterations, the underlying true model is a graph, and CAPRI should be the tool of choice.

Whatever a model is, TRONCO implements a set of functions to assess its confidence via (*i*) non-parametric, (*ii*) parametric and (*iii*) statistical bootstrap.

## 3 Discussion

TRONCO provides up-to-date, theoretically well-founded, statistical methods to understand the evolution of a cancer (ensamble-level) or a single tumor (individual-level). The implemented algorithms are demonstrably the state-of-the-art for the progression inference problem, in terms of computational cost, scalability with respect to sample size, accuracy and robustness against noise in the data. The implementation makes straightforward the interaction of TRONCO with other common bioinformatics tools, possibly allowing the creation of a common suite of tools for cancer progression inference.

Finally, we refer to [3] or the Supplementaty materials for a demonstration of the usage of TRONCO on real genomics data both at the ensemble-level and individual-level progression models. In particular, this paper outlines the capability of the methods to reproduce much of the current knowledge on the progression for a set of cancer types, as well as to suggest clinically relevant insights. Furthermore, we also provide users with detailed manuals, vignettes, and source code to replicate all the analysis presented in the paper plus others (case studies: colorectal cancer, clear cell renal cell carcinoma and acute chronic myeloid leukaemia) in the Supplementary Materials and at the TRONCO official webpage (Software section at http://bimib.disco.unimib.it/).

**Financial support**. MA, GM, GC, AG, DR acknowledge Regione Lombardia (Italy) for the research projects RetroNet through the ASTIL Program [12-4-5148000-40]; U.A 053 and Network Enabled Drug Design project [ID14546A Rif SAL-7], Fondo Accordi Istituzionali 2009. BM acknowledges founding by the NSF grants CCF-0836649, CCF-0926166 and a NCI-PSOC grant.

